# Unmet Expectations About Material Properties Delay Perceptual Decisions

**DOI:** 10.1101/2022.07.28.501825

**Authors:** Amna Malik, Katja Doerschner, Huseyin Boyaci

## Abstract

Based on our expectations about material properties we can implicitly predict an object’s future states, e.g. a wine glass falling down will break when it hits the ground. How these expectations affect relatively low level perceptual decisions, however, has not been systematically studied previously. To seek an answer to this question we conducted a behavioral experiment using animations of various familiar objects (e.g. key, wine glass etc.) freely falling and hitting the ground. During a training session participants first built expectations about the dynamic properties of those objects. Half of the participants (N=28) built expectations consistent with our daily lives (e.g. a key bounces rigidly), whereas the other half learned an anomalous behavior (e.g. a key wobbles). This was followed by experimental sessions, in which expectations were unmet in 20% of the trials. In both training and experimental sessions, participants’ task was to report whether the objects broke or not upon hitting the ground. Critically a specific object always remained intact or broke, only the manner with which it did so differed. For example, a key could wobble or remain rigid, but it never broke. We found that participants’ reaction times were longer when expectations were unmet even when those expectations were anomalous and learned during the training session. Furthermore, we found an interplay between long-term and newly learned expectations, which could be predicted by a Bayesian updating approach. Overall, our results show that expectations about material properties can have an impact on relatively low-level perceptual decision making processes.

## Introduction

Objects are made of certain materials that determine their physical properties. Through the lifetime of experiences, our brain forms long term expectations about the associations between objects and these physical properties (Buckingham, Cant, & Goodale, 2009; Fleming, Wiebel, & Gegenfurtner, 2013). Based on these learned associations, we can predict future states of objects under different forces (Alley, Schmid, & Doerschner, 2020). For instance, when we hold a tea cup in our hand we will be careful not to drop it because we can predict what happens if it falls to the ground. On the other hand, we would not worry a lot if we slip a piece of cloth from the grip of our hand. These expectations are believed to influence behavior through top-down processes and they may often be implicit (Alley et al., 2020; Kersten, Mamassian, & Yuille, 2004; Kveraga, Avniel, & Bar, 2007). Indeed we become aware of our expectations only when we encounter a situation in which they are unmet, or violated, as shown in Figure 1.

**Figure 1:**
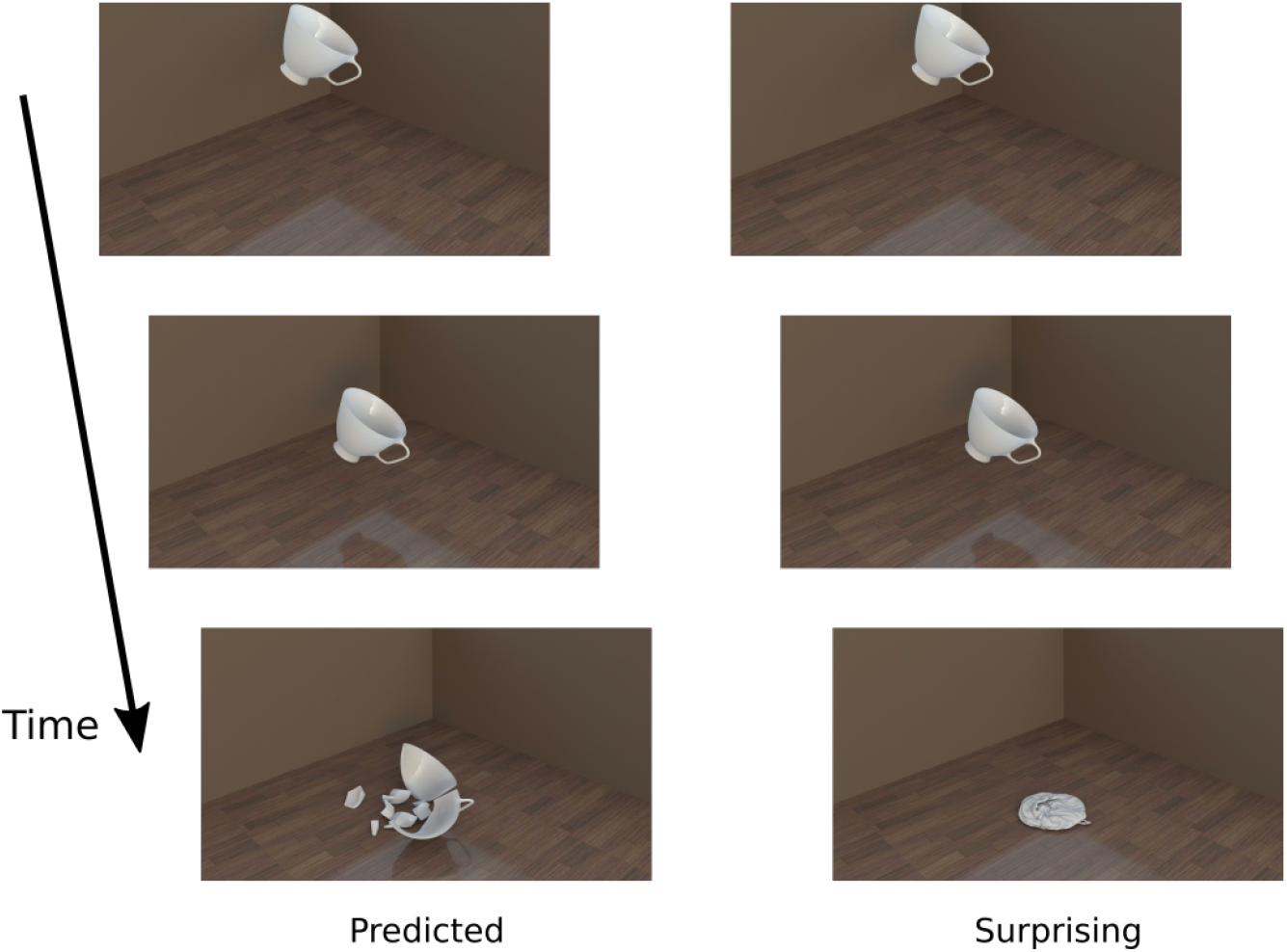
As soon as you see a teacup start falling down, your visual system predicts the future. If not caught, the cup will hit the ground and shatter. But instead if the cup unnaturally wrinkles as a piece of cloth upon hitting the ground, you get surprised and even amazed. Because here your expectations and the visual input mismatch (Alley et al., 2020).

Here we study the effect of long term and newly acquired, context-dependent expectations about material properties on the speed of relatively low level perceptual decisions. A great number of studies have shown that observers perceive the expected stimuli faster (Stein & Peelen, 2015; Summerfield & de Lange, 2014; Wyart, Nobre, & Summerfield, 2012). Those studies, however, usually focused on identification of static stimuli. Only a few studies tested the effects of expectations about material properties in dynamic scenes (Alley et al., 2020). In their study Alley et al. presented participants computer animations of objects that are falling down and behaving in a predicted or surprising way upon hitting the ground. For example a teacup could shatter as predicted or, surprisingly, wrinkle as a piece of cloth. The task of the observers was to judge as quickly and as accurately as possible one of four high-level attributes of the objects in each trial, which were hardness, gelatinousness, heaviness, and liquidity. Alley et al. found that long term expectations bias the perception of high-level material attributes of familiar objects. For example, a spoon that wrinkles upon impact is judged harder than a piece of cloth that also wrinkles. Further, they showed that the reaction times were longer in the surprising trials. In the current study we use a paradigm similar to that in Alley et al. We present the observers computer animations of familiar objects falling down and behaving in a predicted or surprising way upon hitting the ground. To target relatively low level perceptual decisions, however, we do not ask the observers to make judgments about high-level material attributes. Instead, our question is simply “did the object break?” Importantly, breaking objects always break and non-breaking objects always remain intact upon hitting the ground in both expected and surprising conditions. Thus, the correct response for the same object, whether surprising or predicted, does not change, eliminating a response preparation confound. With this paradigm and through measuring the reaction times (RTs), we are able to assess the effect of expectations about material properties on relatively low level perceptual decisions on motion patterns.

In short, our research question is whether expectations about material properties affect low level perceptual decisions. We hypothesize that if they do, then RTs should be different under the predicted and surprising conditions (pre-planned test). To anticipate, under two different experimental manipulations and with two groups of participants, we found that RTs are indeed longer for the surprising trials. Further, we found an interesting interplay between long term expectations and context-dependent regularities, for which we propose possible explanations.

## Materials and Methods

### Participants

Twenty eight participants participated in the experiment. All had normal or corrected to normal vision, and were naive to the purposes of the experiment. Participants gave their written informed consent before the first experimental session, in line with the guidelines by the Declaration of Helsinki. Experimental protocols and procedures were approved by the Research Ethics Committee of Bilkent University, Turkey.

### Stimuli presentation

An LCD color reference monitor (Eizo CG2730, 27 inches, 2560 × 1440 resolution, refresh rate 60 Hz) was used for stimulus presentation. The monitor was the only source of light in an otherwise completely dark room, where the experiment took place. Participants sat on a chair and viewed the monitor from a distance of 60 cm. A chin rest was used to minimize the head movements. Experimental paradigm was programmed with Psychtoolbox on MATLAB, version 2018a (Brainard, 1997).

Stimuli were generated by a professional graphic artist (Aleksa Radakovic) using Cinema 4d, and consisted of computer animations of six objects that act in a certain way when dropped on the ground; three of them break upon hitting the ground (breaking objects: wine glass, pot and teacup) and the other three do not break (non-breaking objects: spoon, key and rod). Each animation consisted of 46 frames. There were two animations for each object. In one set of animations objects behaved in a natural way upon hitting the ground. Specifically, breaking objects shattered and non-breaking objects bounced rigidly after they hit the ground. In the other set, objects behaved in an anomalous way upon hitting the ground: breaking objects graveled, non-breaking objects wobbled. Figure 2 shows examples of these natural and anomalous behaviors.

**Figure 2:**
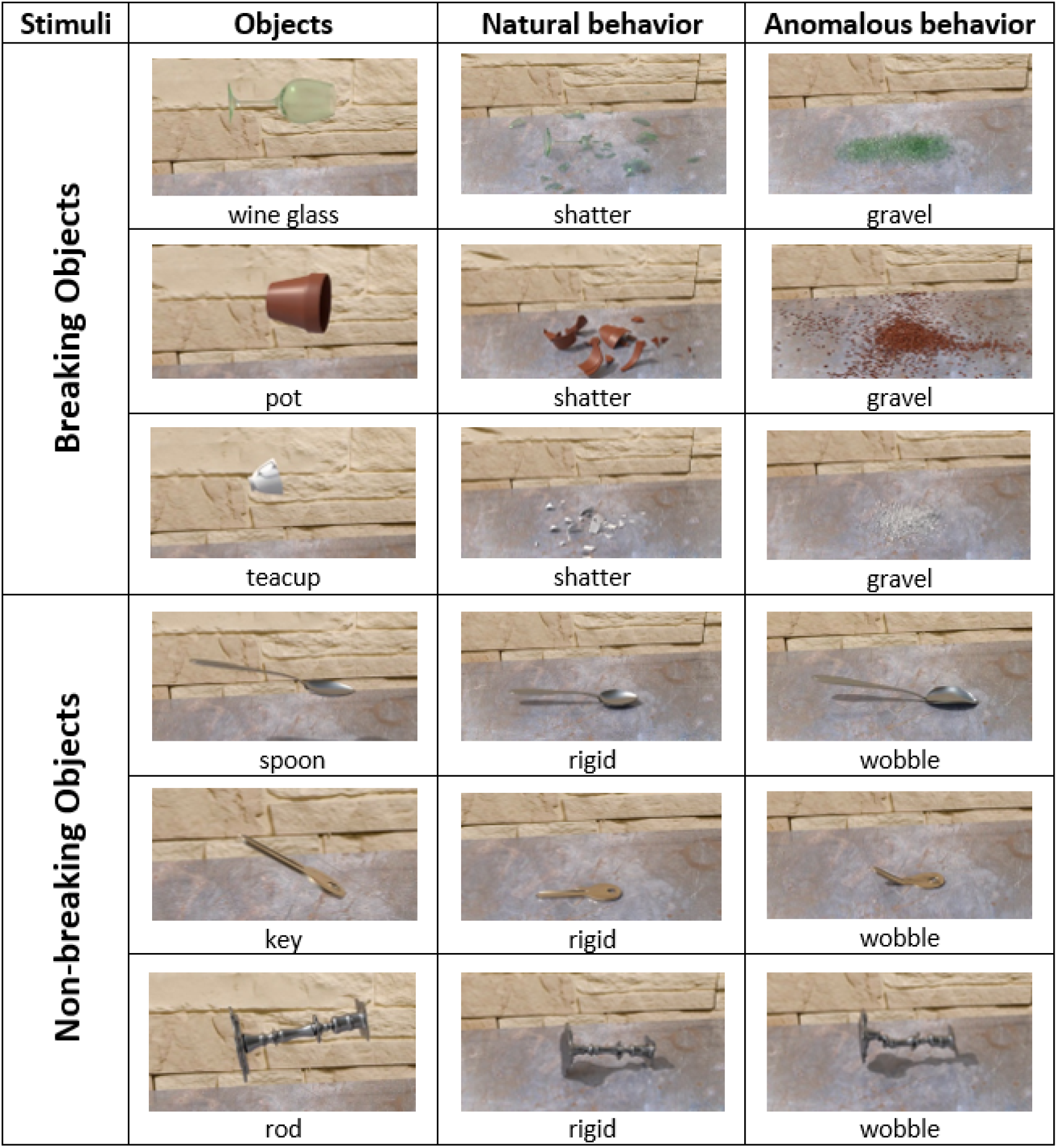
Six objects used as stimuli and their natural and anomalous behaviors. For the participants in group-1 natural behavior was predicted, anomalous behavior was surprising. For the participants in group-2 anomalous behavior was predicted, natural behavior was surprising. These expectations were formed through a training session before the main experiment.

### Experimental Design

Participants were divided into two groups. Each participant underwent a training session followed by an experimental session. During the training session, participants in group-1 were presented with animations where objects behaved naturally, whereas participants in group-2 were presented with animations where objects behaved anomalously (20 trials for each object). Thus, the context-dependent expectations formed for group-2 was different than long-term expectations. During the experimental session 10 animations were shown for each object. Of those 10, 8 were the same as in the training session (for group-1 natural, for group-2 anomalous behavior). We call these predicted trials. The remaining 2 trials were from the untrained category (for group-1 anomalous, for group-2 natural behavior). We call these surprising trials. Order of presentation was randomized in all sessions.

All sessions started with an instruction screen, followed by the animations as soon as any key is pressed. Animations were preceded by a 1-second blank screen with a central fixation cross. Each animation was 1.53 seconds long (46 frames, 30 frames per second). The task was to answer the question, “Did the object break?” by pressing the corresponding keys for “yes” and “no” on the keyboard, after the object hits the ground. In the training session, an error sound was delivered if the participant answered the question before the object hit the ground. Reaction times were measured from the time object makes an impact on the ground (15th frame) to the time when the participant pressed a key. The next trial did not start until the observer responded.

### Analysis

Analyses were performed on MATLAB and JASP (JASP Team, 2022). For both the training and experimental sessions, data from trials in which reaction times are negative (a response made before the object hits the ground), or do not fall within the ±3SD of the mean were excluded from analyses (a total of 24 out of 1680 data points excluded). For the training sessions, reaction times were analyzed using repeated measures ANOVA with “trial numbers” as the repeating factor and “group” as between subject factor. For the experimental sessions, reaction times of predicted trials for breaking and non breaking objects were averaged separately for each participant, and compared to the reaction times of surprising trials with a 3-way repeated measures ANOVA with objects (breaking and non-breaking) and conditions (predicted and surprising) as repeating factors and group as between subject factor. To answer our main research question, namely whether reaction times are longer for surprising trials compared to predicted trials, we performed pre-planned Welch’s tests. In these tests we compared the overall mean reaction times of predicted and surprising trials per group. Further, we compared the mean reaction times for predicted and surprising conditions per participant with paired sample Student’s *t*-tests. Finally, to investigate the effect of expectations specifically on breaking and non-breaking objects, we averaged reaction times separately for breaking and non breaking objects per participant. Then, we compared the mean reaction times of predicted and surprising conditions with a paired sample *t*-test (predicted breaking versus surprising breaking and predicted non-breaking versus surprising non-breaking).

## Results

Figure 3 shows the mean reaction times (RTs) as a variable of trial number in the training session. Repeated measure ANOVA indicated a significant main effect of trial number (*p <* 0.001) and an interaction between groups and trial numbers (*p <* 0.022). Inspecting the plot reveal that RTs get shorter towards the end of the session, which shows that the training was effective. Moreover, Figure 3 shows that the training effect was stronger for participants in group-2. Mean RTs of group-2 were higher than group-1 in the beginning, but they reached group-1 levels and even became slightly shorter after several tens of trials and remained that way until the end of the session.

**Figure 3:**
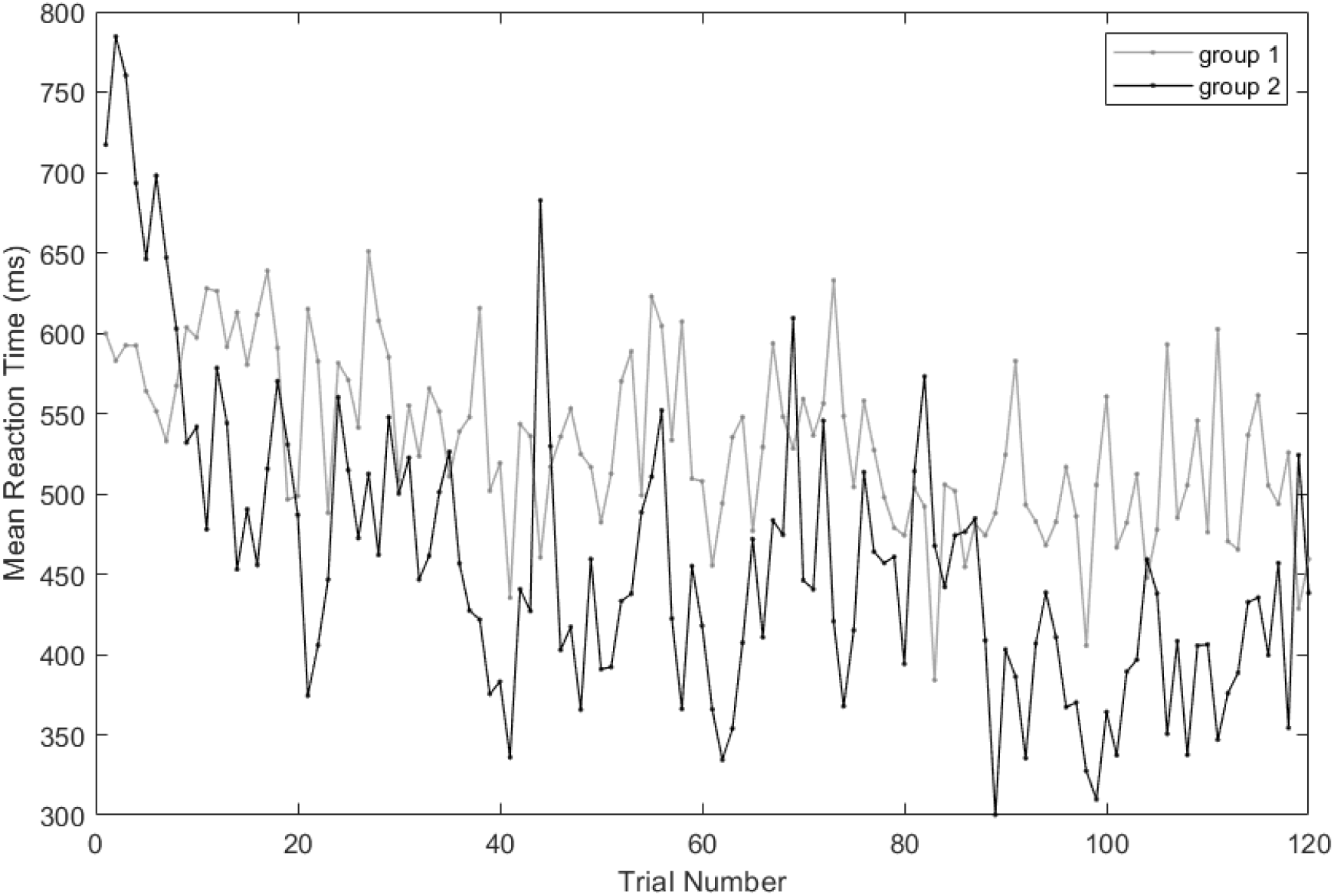
Reaction times (RTs) from the training session. RTs are averaged across participants at each trial number. Clearly the RTs get shorter as the session progresses. This training effect is stronger for the participants in group-2, who were trained on the anomalous behavior.

For the experimental session, a *t*-test showed no effect of expectations (*p* = 0.875) on the percentage of correct responses with a mean of 98.6% for predicted and 98.5% for surprising stimuli, which was anticipated because this is a relatively easy task where breaking objects always break, intact objects always remain intact. Results of 3-way repeated measures ANOVA on RTs, on the other hand, show a significant main effect of conditions (predicted vs surprising, *p <* 0.001), a significant main effect of objects (breaking vs non-breaking, *p <* 0.05), and a significant interaction between objects and groups (*p <* 0.01), between objects and conditions (*p <* 0.01) and between objects, groups and conditions (*p <* 0.001). Table 1 reports the detailed results of the ANOVA.

**Table 1:**
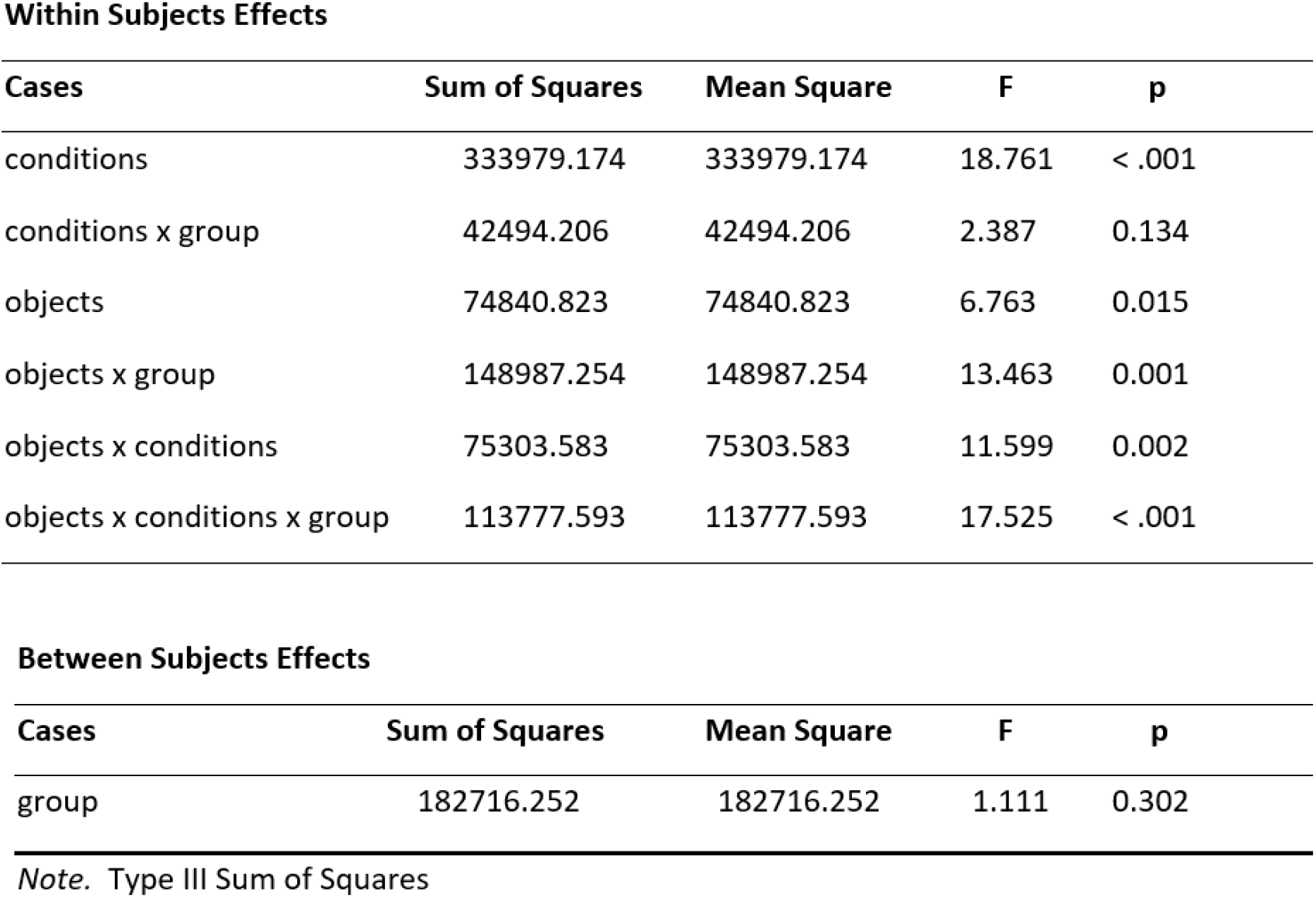
Table showing results 3 way repeated measures ANOVA with objects (breaking and non-breaking) and conditions (predicted and surprising) as repeating factors and group as between subject factor.

Within Subjects Effects
| Cases | Sum of Squares | Mean Square | F | p |
| --- | --- | --- | --- | --- |
| conditions | 333979.174 | 333979.174 | 18.761 | < .001 |
| conditions x group | 42494.206 | 42494.206 | 2.387 | 0.134 |
| objects | 74840.823 | 74840.823 | 6.763 | 0.015 |
| objects x group | 148987.254 | 148987.254 | 13.463 | 0.001 |
| objects x conditions | 75303.583 | 75303.583 | 11.599 | 0.002 |
| objects x conditions x group | 113777.593 | 113777.593 | 17.525 | < .001 |

| Cases | Sum of Squares | Mean Square | F | p |
| --- | --- | --- | --- | --- |
| group | 182716.252 | 182716.252 | 1.111 | 0.302 |

*Note.* Type III Sum of Squares

After showing the robustness of main effects and interactions with ANOVA, we next performed pre-planned tests to answer our main research question, namely whether unmet expectations delay perceptual decisions. Figure 4 shows the mean RTs for predicted and surprising conditions. There was a significant difference between predicted and surprising conditions for both groups (Welch test, *p <* 0.001 for group 1, *p <* 0.01 for group 2). Figure 5 shows the RTs per participant. These results suggest that observers in both groups take longer to respond under the surprising condition. Note, however, that this effect tends to be stronger in group 1.

**Figure 4:**
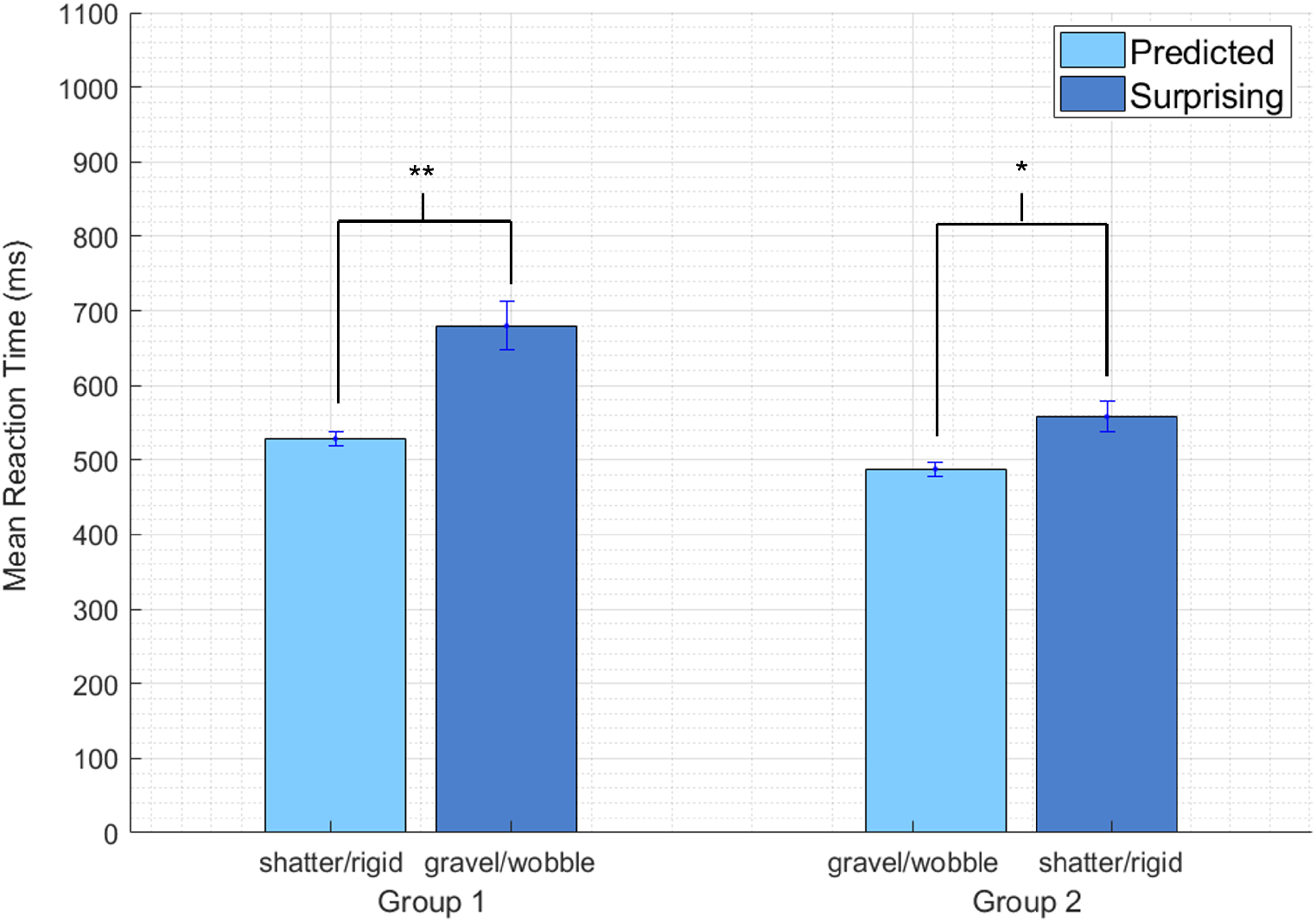
Mean RTs of predicted and surprising conditions averaged across participants (**: *p <* 0.001; *: *p <* 0.01; error bars: SEM.) When expectations are unmet, whether natural or anomalous, perceptual decisions are delayed.

**Figure 5:**
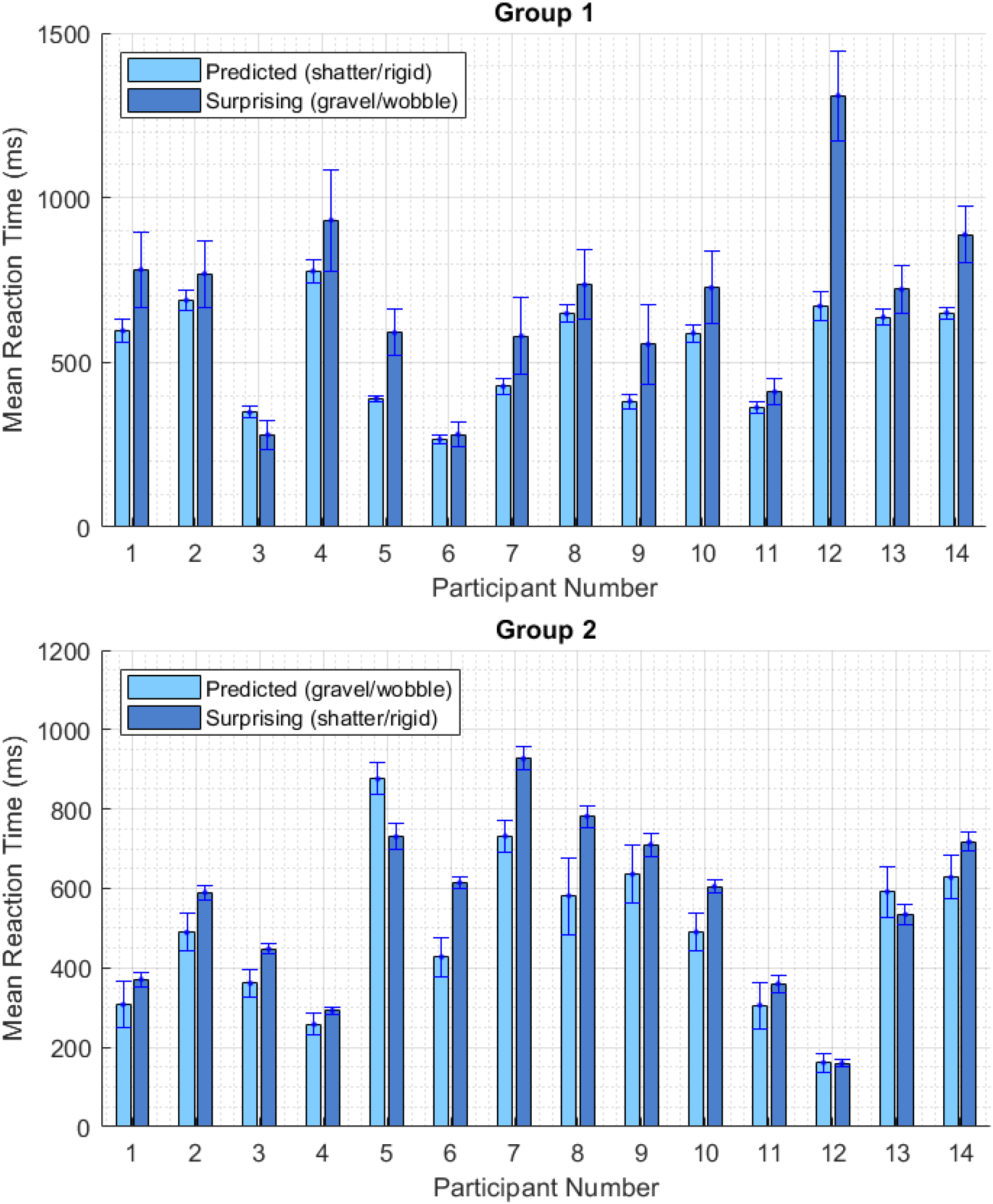
Mean RTs per participant.

Noting that the effect tends to be stronger in group 1, we performed further analyses. For this, we compared the RTs for breaking an non-breaking objects separately. Figure 6 shows those RTs. For group 1, the difference between the RTs under the predicted and surprising conditions was significant for the non-breaking objects but not for the breaking objects. For group 2 the situation was reversed: the difference between the RTs under the predicted and surprising conditions was significant for the breaking objects but not for the non-breaking objects (*p <* 0.0125, corrected for multiple comparisons). We discuss this finding below.

**Figure 6:**
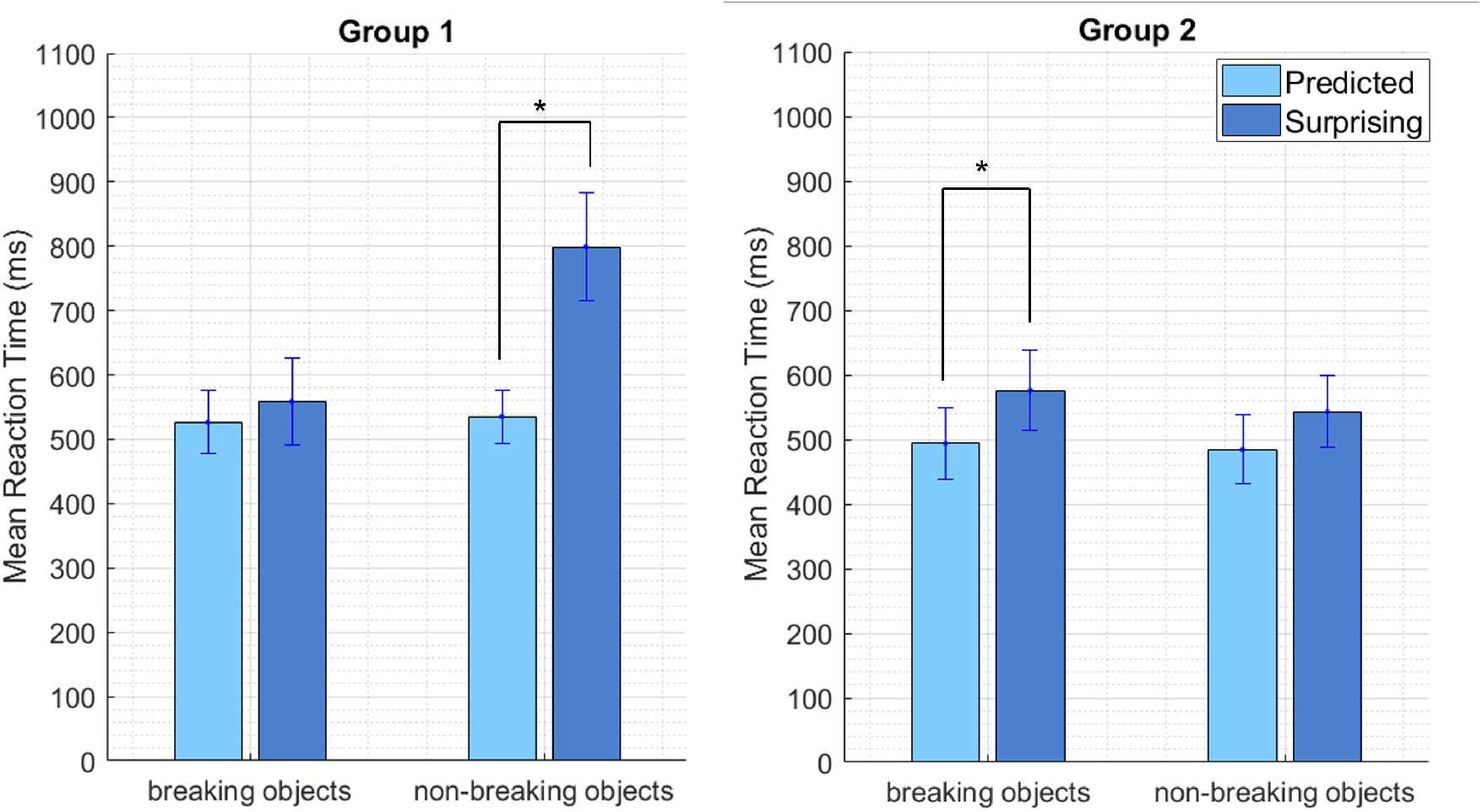
Mean RTs for breaking and non breaking objects plotted separately (*: *p <* 0.0125, corrected for multiple comparisons)

## Discussion

Here we studied the effect of expectations about material properties on the speed of relatively low-level perceptual decisions. We presented computer animations of objects falling down, and asked the participants to report as soon as possible whether the objects break or not upon hitting the ground. We found that participants were slower to make this judgment when their expectations about the material properties were not met. Furthermore, this was true even when participants were trained to predict an anomalous behavior, for example a candle stick to bounce as if made of jelly. The pattern of our results can not be explained by motor response preparations, because whether under the predicted or surprising condition, for a given object the correct response was always the same: breaking objects always broke, intact objects always remained intact. Motion statistics, on the other hand, might have affected the RTs. For example it could be easier to decide that an object remains intact with the motion statistics of a rigid body compared to a gelatinous one. Those low level motion statistics, however, cannot explain the differences between the two experimental groups. Thus, our results show that unmet expectations about material properties delay perceptual decisions.

### Expectations about material properties affect low-level perceptual processes

Our results show that when expectations are not met perceptual decisions are delayed. This is in line with previous studies. For example Alley et al. (2020) found that unmet expectations delay participants’ decisions about material attributes. But unlike in most previous literature, in our study participants’ task was not about material attributes. Thus they did not need to attend and process the material properties, they only needed to analyze the motion patterns after the objects hit the ground. A sensible strategy could be to ignore the object-material associations, and focus entirely on the low-level motion patterns after the impact. Nevertheless, participants’ expectations about material properties still affected the speed of their decisions. These results demonstrate that high-level expectations can affect low-level perceptual processes, even when those expectations are task irrelevant.

### Training alters expectations

Our daily subjective experiences suggest that humans have long term expectations about object-material associations, and static and dynamic properties of materials. Systematic studies in literature have shown that observers use a variety of visual cues to estimate material properties of objects. For example, observers use shape, optic and motion cues to judge the stiffness of materials (Doerschner et al., 2011; Paulun, Schmidt, Assen, & Fleming, 2017; Schmidt, Paulun, Assen, & Fleming, 2017). These associations not only help us to recognize and identify the object and materials efficiently but also help in action planning and guiding our interaction with them (Buckingham et al., 2009; Doerschner et al., 2011; Sutter, Drewing, & Müsseler, 2014)

Some long term expectations are “stubborn” and do not easily change, but some can be altered under experimental conditions (de Lange, Heilbron, & Kok, 2018; Yon, de Lange, & Press, 2019). For example, Adams, Graf, and Ernst (2004) showed that “light from above” prior could be altered when participants are trained with haptic feedback. Similarly, Sotiropoulos, Seitz, and Seriès (2011) showed that “slow speed prior”, which explains many motion and direction illusions, can be altered through training sessions. The pattern of RTs we found in the current study is consistent with this literature. We found that RTs of group 2 were longer under the surprising condition compared to the predicted condition, even though the predicted anomalous behaviors were in conflict with the long term expectations. This shows that participants learned new context-dependent expectations during the training session.

RT data from the training session provides further insights about the progress of learning. Firstly, the decrease in RTs was larger for group 2 compared to group 1. This was anticipated because only in group 2 participants learned new associations and formed new context-dependent expectations. In the beginning of the training sessions RTs of group 2 were longer than those of group 1, which is also anticipated because the object behaviors were anomalous and not predicted based on long term expectations. But as the session progressed the group 2 participants started to learn to expect an anomalous behavior in the context of the experiment, and their RTs decreased. Towards the end of the session RTs of group 2 were equal to, and even slightly lower than RTs of group 1. This further reduction might be related to an ‘oops’ factor, whereby a sequence of asynchronously presented mismatching cues can lead to efficient learning (Adams, Kerrigan, & Graf, 2010).

### Interplay between long term expectations and context-dependent regularities

The difference between the RTs under the predicted and surprising conditions tended to be larger for group 1 compared to group 2. Thus, the overall effect of expectations tended to be stronger in group 1 compared to group 2. For group 1, where long term expectations and context-dependent regularities were consistent, a strong expectation effect was indeed anticipated. Whereas for group 2 long-term expectations, which can often be strong (Seriès & Seitz, 2013), conflicted the context-dependent regularities. This conflict could have reduced the overall strength of the newly acquired context-dependent expectations in group 2. Further scrutiny revealed a significant effect of expectation for intact objects but not for breaking objects in group 1. Conversely, for group 2 there was a significant effect for breaking objects but not intact objects. This finding might seem puzzling at first but it can be explained by different strengths of long term expectations. Long term expectations for the non-breaking objects used in the experiment, such as the candle-stick, to be rigid rather than gelatinous might be very strong, leading to the significant effect found for those objects in group 1. These long term expectations, however, strongly conflict the context-dependent regularities for group 2, and thus produce weaker new expectations and result in no effect for the non-breaking objects in that group. Conversely, for the breaking objects used in the experiment, the long term expectations to shatter might not be that strong, leading to little or no effect of expectation in group 1. But this time, because the long term expectations are weak, the newly-acquired expectations are stronger and this results in a significant effect for group 2.

### Bayesian updating

In this part we discuss a Bayesian updating approach that can formally explain the pattern of our findings. In its basic form, Bayesian rule allows computing the posterior distribution of the world states given the observation by simply combining the prior probability distribution of the world states (*i*.*e*. the expectations) and the likelihood function of those world states under the observed data. This process can be dynamic, for example the posterior computed at one moment can be used as the prior of the next (Bitzer, Park, Blankenburg, & Kiebel, 2014; Urgen & Boyaci, 2021). This is called Bayesian updating of the posterior. The conceptual ideas provided above to explain our findings can be formulated in such an updating model. In such a model, computations in a trial would continue until enough evidence is collected to reach a decision, which is breaking versus non-breaking in our experiment. In case initial prior and the likelihood agree, this computation can reach a decision relatively quickly, because the posterior distribution would be sharp and clearly favor one of the world states. Whereas if the prior and likelihood disagree, posterior distribution would become broader making it harder to make a decision, and the computation would need to continue. For example, for group 2 in the beginning of the training session the prior distribution and the likelihood function largely disagree, thus the model would predict longer RTs consistent with the empirical data. But as the priors are updated, the discrepancy between them reduces, thus in later trials computations would converge quicker and the model would predict shorter RTs, again consistent with the empirical data. The same logic applies to the trials in the experimental session. In short, a Bayesian updating approach can formally explain the empirical findings of the current study.

## Conclusion

To conclude, we found that unmet expectations about dynamic material properties delay perceptual decisions. We argue that high-level expectations about material properties affect relatively low-level perceptual processes even when those expectations are not directly task-relevant. Furthermore, we show that through training participants form new context-dependent expectations. Those newly formed context-dependent expectations and long term expectations together shape the perceptual processes, which can be formulated using a Bayesian updating approach.

## References

Adams, W. J., Graf, E. W., & Ernst, M. O. (2004). Experience can change the ‘light-from-above’ prior. Nature Neuroscience, 7 (10), 1057–1058. doi: 10.1038/nn1312

Adams, W. J., Kerrigan, I. S., & Graf, E. W. (2010). Efficient visual recalibration from either visual or haptic feedback: the importance of being wrong. Journal of Neuroscience, 30 (44), 14745–14749.

Alley, L. M., Schmid, A. C., & Doerschner, K. (2020). Expectations affect the perception of material properties. Journal of Vision, 20 (12), 1. doi: 10.1167/jov.20.12.1

Bitzer, S., Park, H., Blankenburg, F., & Kiebel, S. J. (2014). Perceptual decision making: drift-diffusion model is equivalent to a Bayesian model. Frontiers in human neuroscience, 8 (February), 102:1–17. doi: 10.3389/fnhum.2014.00102

Brainard, D. H. (1997). The Psychophysics Toolbox. Spatial Vision, 10, 433–436.

Buckingham, G., Cant, J., & Goodale, M. (2009). Living in a material world: How visual cues to material properties affect the way that we lift objects and perceive their weight. Journal of neurophysiology, 102, 3111–8. doi: 10.1152/jn.00515.2009

de Lange, F. P., Heilbron, M., & Kok, P. (2018). How Do Expectations Shape Perception? Trends in Cognitive Sciences, 22 (9), 764–779. doi: 10.1016/j.tics.2018.06.002

Doerschner, K., Fleming, R. W., Yilmaz, O., Schrater, P. R., Hartung, B., & Kersten, D. (2011). Visual motion and the perception of surface material. Current Biology, 21 (23), 2010–2016.

Fleming, R., Wiebel, C., & Gegenfurtner, K. (2013). Perceptual qualities and material classes. Journal of Vision, 13. doi: 10.1167/13.8.9

JASP Team. (2022). JASP (Version 0.16.3)[Computer software]. Retrieved from https://jasp-stats.org/

Kersten, D., Mamassian, P., & Yuille, A. (2004). Object perception as bayesian inference. Annual review of psychology, 55, 271–304. doi: 10.1146/annurev.psych.55.090902.142005

Kveraga, K., Avniel, G., & Bar, M. (2007). Top-down predictions in the cognitive brain. Brain and cognition, 65, 145–68. doi: 10.1016/j.bandc.2007.06.007

Paulun, V., Schmidt, F., Assen, J., & Fleming, R. (2017). Shape, motion, and optical cues to stiffness of elastic objects. Journal of Vision, 17(1), 1–22. doi: 10.1167/17.1.20

Schmidt, F., Paulun, V., Assen, J., & Fleming, R. (2017). Inferring the stiffness of unfamiliar objects from optical, shape, and motion cues. Journal of Vision, 17(3), 1–17. doi: 10.1167/17.3.18

Seriès, P., & Seitz, A. (2013). Learning what to expect (in visual perception). Frontiers in human neuroscience, 7, 668.

Sotiropoulos, G., Seitz, A. R., & Seriès, P. (2011). Changing expectations about speed alters perceived motion direction. Current Biology, 21 (21), R883–R884.

Stein, T., & Peelen, M. (2015). Content-specific expectations enhance stimulus detectability by increasing perceptual sensitivity. Journal of experimental psychology. General, 144. doi: 10.1037/xge0000109

Summerfield, C., & de Lange, F. (2014). Expectation in perceptual decision making: Neural and computational mechanisms. Nature reviews. Neuroscience, 15. doi: 10.1038/nrn3838

Sutter, C., Drewing, K., & Müsseler, J. (2014). Multisensory integration in action control (Vol. 5). Frontiers Media SA.

Urgen, B. M., & Boyaci, H. (2021). Unmet expectations delay sensory processes. Vision Research, 181, 1–9. doi: 10.1016/j.visres.2020.12.004

Wyart, V., Nobre, A., & Summerfield, C. (2012). Dissociable prior influences of signal probability and relevance on visual contrast sensitivity. Proceedings of the National Academy of Sciences of the United States of America, 109, 3593–8. doi: 10.1073/pnas.1120118109

Yon, D., de Lange, F. P., & Press, C. (2019). The Predictive Brain as a Stubborn Scientist. Trends in Cognitive Sciences, 23 (1), 6–8. doi: 10.1016/j.tics.2018.10.003

